# Multi-Layer Brain-Mimicking Phantom for Replicating Dura and Pia Membrane Dimpling and Rupture Properties During Neural Interface Implantation

**DOI:** 10.64898/2026.02.05.704082

**Authors:** Dongyang Yi, Kevin Lat, Lei Chen

## Abstract

Development of novel neural interfaces faces buckling challenges and heavily relies on trial-and-error tests via *in vivo* animal brain insertions for design optimizations towards the minimal-damaging version for enhanced recording and stimulation outcome. To enable low-cost and fast-turnaround neural interface development and to enable previously impossible insertions via new understanding of the cutting process, this study developed a reproducible, multi-layer brain-mimicking phantom designed to replicate the rodent pia and dura mater dimpling and rupture force performance observed during *in vivo* tests. The phantom was composed of a 0.5% (w/v) agarose cortex layer, a 1.01% (w/v) agarose pia mater layer, and a pre-stretched polyvinyl chloride (PVC) dura mater layer, assembled via easily duplicable benchtop protocols. Using a cantilever-beam force measurement system, rupture force and dimpling depth were quantified across microwires of varying diameters (12–100 μm), materials (tungsten, stainless steel), and tip geometries, as well as segmented silicon probe shanks. Phantom test results closely matched *in vivo* Sprague–Dawley rat data, validating the performance of the developed multi-layer phantom. At the same time, phantom insertion trial variability was substantially lower than *in vivo* tests, enabling a repeatable, low-cost, early-stage screening platform of novel electrode designs. The phantom’s modular design also allowed tuning of layer thickness and stifness of each layer for diferent species or devices, ofering a validated customizable testing platform to accelerate novel neural implant development and reduce animal use.

## 1. Introduction

Capturing single-neuron activity is essential for decoding the complex, interconnected functions of the brain that underline cognition, behavior, and disease mechanisms [1-4]. *In vivo* measurement with invasive microelectrode arrays (MEAs) remain the gold standard for acquiring such single-neuron signals, as they provide direct access to neuronal activity with high spatial and temporal resolutions [5-8]. However, to fully understand the broad spectrum of brain function, there is a growing need for implantable, high-channel-count MEAs capable of chronic recordings across multiple brain regions [9, 10]. Meeting these demands has driven significant efforts to develop smaller, more flexible microelectrodes that minimize tissue damage while maintaining long-term functionality. Despite these advancements, the transition to smaller and more flexible electrodes (e.g., cellular-scale microwire electrodes and flexible-substrate probes) has introduced new challenges, particularly in terms of electrode surgical implantation through the membranes (e.g., the dura and pia maters) [11, 12]. When pushed against the membranes during brain insertion, the small and flexible electrodes are susceptible to structural instability, causing buckling and/or severe deflection, which can prevent proper targeting and degrade recording fidelity and must be addressed to enable reliable and widespread use of such miniaturized chronic neural interfaces [5, 13].

To address the buckling challenge, surgical protocol and ad hoc design modifications such as tip sharpening, insertion shuttles, temporary resolvable support, and incremental insertion have been employed by researchers to strengthen the electrodes during implantation [14-22]. While all the strategies provided improvement on the insertion process, the design optimization (e.g., best sharp tip angle, minimal shuttle size) for novel neural interface development still heavily rely on trial-and-error tests via *in vivo* animal brain insertions. While the *in vivo* tests do stand as the gold standard for verification, the high-cost, labor-intensiveness, and natural variance involved in animal tests could significantly slow down the preliminary iterative development and optimization of novel neural interfaces and the corresponding implantation strategies. Such surgical animal trial needs also prevent thorough understanding of the neural interface implantation mechanics involving electrode-membrane interactions, leading to unnecessarily large, stiff, and damaging neural interface and related implantation instruments to be used. Also, the thick and tough dura mater layer is usually treated as impossible to penetrate and thus surgically removed, causing extra trauma and preventing brain-wide electrophysiological implantations and recording/stimulation.

To avoid large number of animals for low-cost and fast-turnaround neural interface development and to enable previously impossible insertions via new understanding of the cutting process, brain-mimicking phantoms, which serve as synthetic surrogates for biological tissues, could serve as a promising alternative [23]. By providing a controlled and repeatable test environment as compared to animals, phantoms can complement *in vivo* studies and accelerate the development of advanced microelectrode technologies. A number of brain phantoms have been proposed: for example, 0.6% agarose gels have effectively mimicked intraparenchymal catheter insertion forces [24, 25], while PVA-cryogels have been used to fabricate anatomically accurate brain structures for multimodal imaging validation [26]. Additional models, such as silicone-based phantoms [27, 28] and complex agarose or gelatin constructs designed for optical or mechanical calibration [29-34], have served roles in surgical training and imaging applications. In neurosurgery, multiple biological and synthetic materials have also been used as the dural substitutes to repair the dura mater, mimicking the long-term biological, electrical, chemical, and mechanical properties of the real brain membranes [31, 35-38].These phantoms are widely used in biomedical research and healthcare due to their ability to replicate specific brain layer properties, cost-effectiveness, ease of preparation, and consistent performance. However, to the best of our knowledge, there lacks a brain-mimicking phantom material that replicates the brain tissue and membrane deformation and rupture properties against neural interface insertions.

Development of such brain phantom faces key difficulties that it is not practical to surgically obtain brain membrane (dura and pia maters) samples in certain size and configuration for standardized mechanical property tests while keeping its *in vivo* material properties and status. Instead, the phantom development will focus on matching key implantation outcome parameters as achieved from *in vivo* insertion tests. From a neural interface implantation perspective, two parameters are most commonly used and related to the electrode penetration of the brain membranes: rupture force and dimpling depth at the electrode penetration point. The electrode will buckle if the maximum insertion force (which usually happens at the tough membrane rupture point as friction along further insertion through soft brain tissue (cortex) are comparatively smaller with less unsupported length) is larger than the critical buckling load. On the other hand, as *in vivo* real-time insertion force measurement is difficult during animal surgery, the brain membrane dimpling depth is commonly used as an intuitive indicator of the insertion resistance [5, 39].

In this study, we are developing the brain mimicking phantom based on rupture force and dimpling depth measurement data from *in vivo* Sprague-Dawley rats insertion tests conducted before [5]. Since the dura mater, pia mater, and soft brain cortex tissue have very different material properties, we present a multilayer brain-mimicking phantom solution to replicate both the pia-only and dura-pia penetration cases during neural interface implantation procedure. Agarose layers of different concentration are used as the pia mater and brain tissue phantom materials and a thin polyvinyl chloride (PVC) film is chosen as the dura mater phantom layer. The methods section elaborates the reproducible benchtop fabrication protocols for the multi-layer phantom as well as the specific formula found to match Sprague-Dawley rats *in vivo* data. The force and dimpling measurement setup and design of experiments to evaluate the developed phantom performance is also presented. The results section compares the phantom insertion force and dimpling outcomes with *in vivo* data with various types of electrodes to validate the developed phantom. Further development and generalization of the multi-layer rodent brain phantom developing methodology to other animal models and insertion devices are then discussed followed by conclusion of the study.

## 2. Methods

### 2.1. Multi-layer brain-mimicking phantom fabrication protocols

To accurately replicate the mechanical properties and layered structure of brain together with the protective membrane layers during neural interface implantation, a multi-layered brain-mimicking phantom was developed. The phantom consists of three distinct layers: (1) the brain tissue (cortex) layer, (2) the pia mater layer, and (3) the dura mater layer. This design reflects the *in vivo* surgical conditions where microelectrodes are inserted into the brain either with the dura layer intact (dura-pia penetration) or removed (pia only penetration). By incorporating these layers, the phantom captures the multi-layer nature of brain tissue, enabling fine tuning of each layer’s property and generating a more realistic simulation of tissue-electrode interactions on the rupture force and surface dimpling at membrane penetration point.

#### 2.1.1. Agarose-based pia-brain phantom layer fabrication

The pia-cortex layers, representing the pia mater and the brain tissue, was fabricated using a custom agarose-based formulation following the procedure as shown in Fig. 1. Agarose has been widely used as brain-mimicking phantom material for its ability to mimic the soft, viscoelastic properties of brain tissue [24, 25, 40]. For the brain tissue (cortex) layer of the phantom, a 0.5% weight by volume (w/v) agarose solution was first prepared by dissolving agarose powder (RPI A20030, RPI Corp., IL, USA) in deionized (DI) water and heating the mixture with a microwave oven until boiling occurs in the solution. During the heating process, the mixture was additionally swirled around in the container every 30 seconds to have the solution homogenize further. After boiling, the solution went through around 5 minutes of cooling and then 10 mL of the agarose solution (controlled by two 5 mL pipette doses) was pipetted into each section of a silicone ice cube tray mold (each cube dimensions: 31.75 mm x 31.75 mm x 31.75 mm) (Fig. 1(a) and (b)), yielding about 9.9 mm thick brain tissue layer thickness. The tray was then placed on a flat benchtop surface for 10 minutes to have the solution leveled and bubbles eliminated (Fig. 1(c)). A dampened piece of paper towel was then used to cover the entire silicone ice cube tray to maintain the moisture of the material. The covered silicone mold tray was then transferred into a refrigerator (4°C) for 1 hour (Fig. 1(d)) to form a gel-like material with mechanical properties similar to those of rat brain tissue.

**Figure 1.**
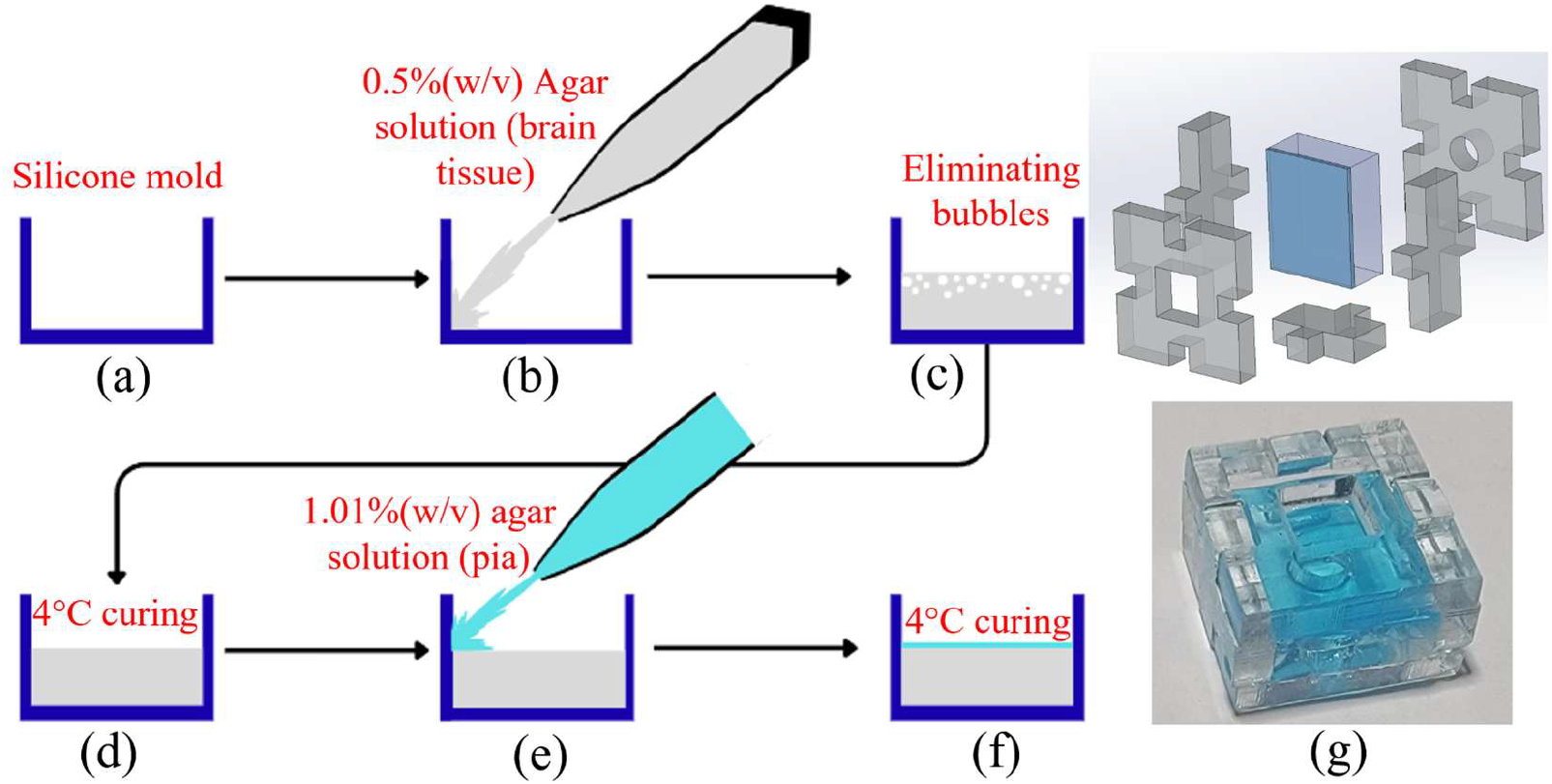
**Fabrication procedure of the dual-layer pia–cortex brain-mimicking phantom: (a) silicone ice cube tray as the phantom mold, (b) pipette injection of brain tissue (cortex) phantom solution into the mold, (c) settling and leveling on a flat benchtop surface while eliminating bubbles, (d) one hour refrigerating for curing, (e) further pipetting of the pia mimicking phantom layer, (f) leveling, bubble elimination, and then curing in refrigerator, and (g) top: custom design and made acrylic holder for the pia-cortex dual-layer phantom and bottom: picture of a fully assembled dual-layer pia-cortex phantom**.

To fabricate the thin, tough pia layer on top of the brain tissue, a 1.01% (w/v) agarose solution was similarly mixed in DI water and brought to a boil. After 3 minutes of cooling, a few drops of food coloring (blue in this study) were added to the pia phantom layer solution for better visibility. 0.6 mL of the dyed solution was pipetted on top of the previously refrigerated phantom (Fig. 1(d)) as a thin film of about 0.6 mm thick above the brain tissue layer surface (Fig. 1(e)). This new pia layer was then covered under a damp paper towel and air cooled for 5 minutes on a flat benchtop surface. The silicone mold with both brain and pia phantom layers was thus leveled during this step to ensure smoothness of the surface. Then, the leveled phantom assembly in the silicone mold was placed back into the refrigerator (4°C) to cool for another hour (Fig. 1(f)). This two-step process ensured a seamless integration of the pia and cortex layers, mimicking the *in vivo* pia-brain structure. For the pia-cortex phantom insertion tests, a custom-designed acrylic frame was used to hold and support the agarose-gel-based phantom layers. The frame was composed of five side panels (Fig. 1(g)) laser-cut from 4.5 mm thick acrylic sheets to ensure dimensional precision and consistency across trials. The internal dimensions of the assembly matched the mold used for gel casting, enabling seamless transfer of the gel from the silicone molding to this test fixture without deformation. The acrylic structure provided a rigid enclosure that maintained the geometric integrity of the phantom during handling and testing. Figure 1(g) illustrates the completed pia-cortex phantom housed within the acrylic assembly, with the upper pia layer visible as a blue, smooth, transparent film atop the brain-mimicking cortex gel at the bottom.

#### 2.1.2. PVC-based dura mater phantom fabrication and assembly

The dura layer, which is notably tougher than the pia and cortex layers, was hard to mimic by the soft Agarose gel. Thus, the dura phantom was fabricated using a polyvinyl chloride (PVC) plastisol formulation (Regular plastic by M-F Manufacturing, Ft. Worth, TX, USA). PVC plastisol polymer solution has been used as tissue mimicking material with tunable mechanical and imaging properties [23]. Its wide range of elastic modulus, hardness, and viscoelastic stress relaxation time constants made it a good choice for the thin but hard to penetrate dura mater layer.

In this study, the PVC mixture used to make dura layer phantom was prepared following the steps as shown in Fig. 2. As in Fig. 2(a-c), PVC polymer solution was mixed with a plastic softener (phthalate ester by M-F Manufacturing, Ft. Worth, TX, USA) in a 2:1 weight ratio. The two components were then mixed together via stirring. Next, the mixed solution was heated until reaching 150°C while keeping stirring and turning clear, forming a flexible rubber-like PVC mixture Fig. 2(d-f).

**Figure 2.**
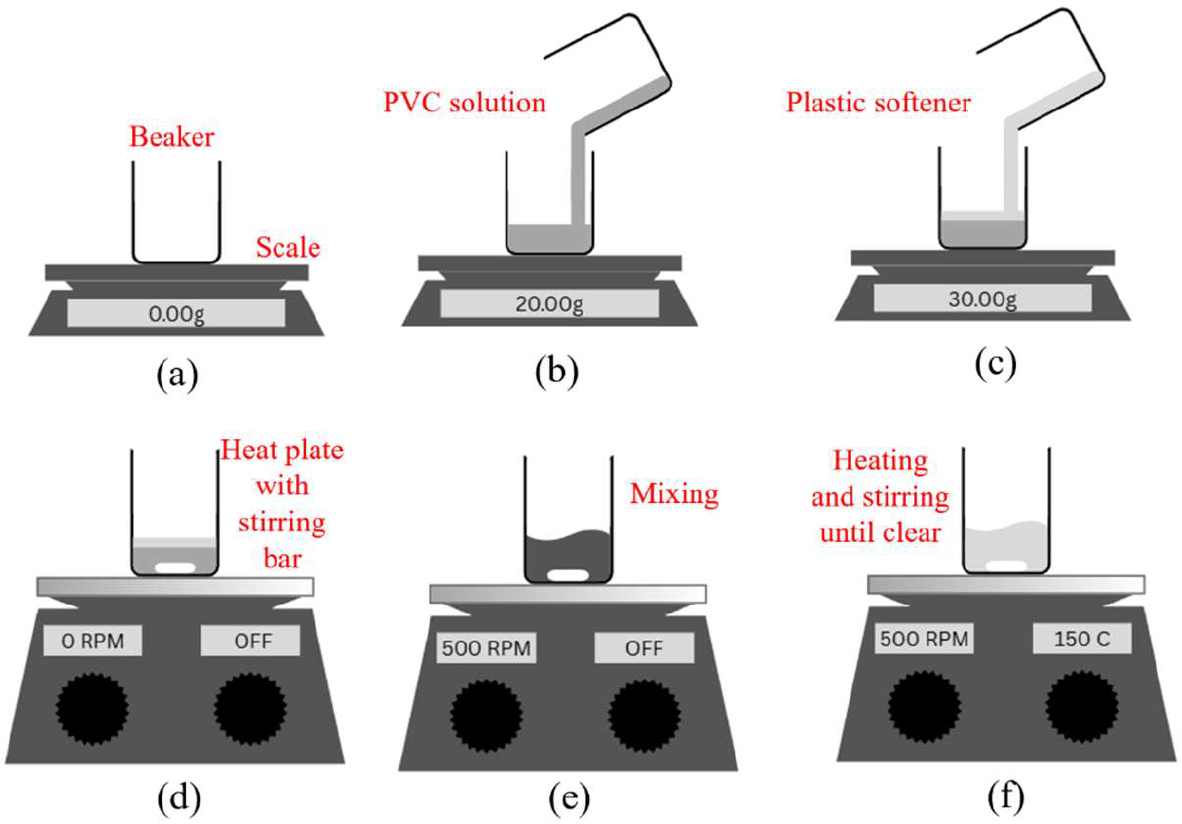
Fabrication process of PVC dura mixture, including mixing, stirring, and heating of liquid PVC solution with plastic softener under a 2:1 weight ratio.

Steps to make the PVC dura mixture into a thin-layer phantom like the dura mater are as elaborated in Fig. 3. To build a custom press mold, two glass cover slips (0.15 mm thick) were placed on both ends of one flat borosilicate glass slide (Fig. 3(a)). A small amount of the dura mater PVC mixture was then poured onto the center of the glass slide, as in Fig. 3(b). Another flat glass slide was placed on top of the first one, thus forming a press mold with the cover slips as spacers and thus the formed PVC thickness was accurately controlled by the cover slip thickness (Fig. 3(c)). Once the thin layer was formed, the assembly was transferred to a refrigerator for 5 minutes to cool and solidify, before retracting the formed thin PVC film from the press (Fig. 3(d)). In this study, a rectangle measuring 25.4 mm by 30 mm was desired for the dura PVC film and it was measured, marked (Fig. 3(e)), and cut using razor blade to the desired shape (Fig. 3(f)).

**Figure 3.**
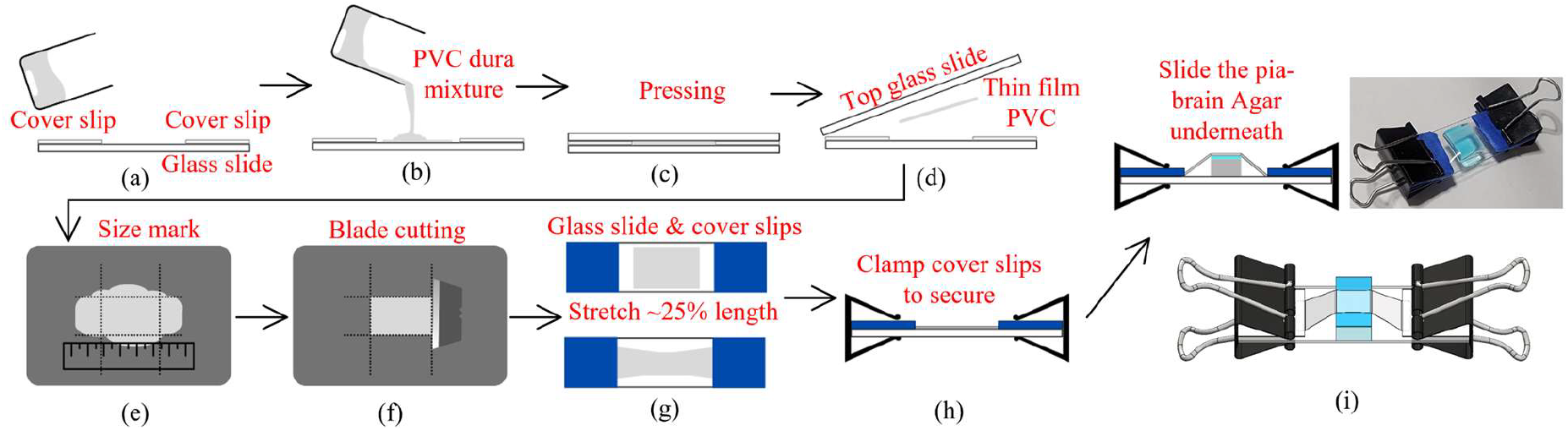
**Fabrication of the PVC-based dura phantom layer and assembly of the multi-layer dura-pia-brain phantom: (a) glass slide and cover slip assembly as the molding base, (b) PVC dura mixture pouring at the glass slide center, (c) pressing from top with another glass slide to create even height PCV film depending on cover slip thickness, (d) thin PVC film de-molding from the assembly after colling and solidification in a refrigerator, (e) mark desired PVC phantom layer size on the achieved film, (f) razor blade trimming to desired size, (g) pre-stretch the PVC film by about 25% length using new glass slide and two cover clips to hold both ends, (h) fixation of stretched PVC phantom by paper clips on both sides, and (i) slide the pre-made agarose-based pia-brain phantom underneath the stretched PVC-based dura layer to achieve the final multi-layer brain phantom**.

Our insertion trials showed that for the dura mater to be in close contact with the pia and brain to provide the pre-rupture resistance (instead of freely sliding and wrinkling on the pia mater surface), it needs to be pre-stretched and held in place. Thus, the cut PVC film was placed on another glass slide with cover slips on both sides. The slips were used to hold the edges of the PVC and stretch them by about 25% of its original length (Fig. 3(g)). After stretching, cover slips on both sides were clamped in placed by paper clips (Fig. 3(h)). To build the dura-pia-brain multi-layer configuration, the pia-brain assembly as shown in Fig. 2 was removed from the acrylic enclosure and slide under the secured and stretched PVC dura layer (Fig. 3(i)). This setup ensured uniform contact and minimized the formation of air pockets or folds between layers. The transparent nature of the dura material and the underlying layers enabled real-time visual confirmation of alignment and contact before testing as well as membrane layer rupture during insertion tests.

### 2.2. Experimental setup for brain-mimicking phantom evaluation

The main objective of this study is to develop a easily duplicable while repeatable multi-layer brain-mimicking phantom system that could match our rupture force and dimpling depth measurement from *in vivo* implantation tests on Sprague-Dawley rat brain [5], under both pia-only and dura-pia insertion conditions, for all different types of microwires and silicon probes tested previously on rats. Thus, this research inherited key components from the *in vivo* study, including the same microwire electrodes with variant tip geometries, same silicon probe shanks, similar force measurement apparatus, and critical parameters for evaluating tissue-electrode interactions (e.g., membrane rupture force and dimpling depth at rupture).

In the *in vivo* study, the small rupture force and dimpling depth were measured by a specially designed flexible cantilever-beam based force measurement system. In this study, to evaluate the developed brain phantom performance, a similar system was developed with minor modifications to accommodate its applicability for phantom-based experiments. As shown in Fig. 4, the force and dimpling measurement setup consists of a cantilever beam, a 3D-printed cantilever beam cap for microwire fixation, and three computer-controlled linear stages (MOX-02-30 by Optics Focus, China) to enable precise movement along the X-, Y-, and Z-axes. A laser displacement sensor (LK-G10 by Keyence, Osaka, Japan) was used to measure the deflection of the cantilever beam during microelectrode insertion. The system was designed to provide high-resolution force, ensuring accurate capture of tissue-electrode interactions. In this study, the cantilever beam was constructed from AISI 304 stainless steel, with dimensions of 40 mm in width, 0.9144 mm in thickness, and 380 mm in length. The laser displacement sensor was positioned at a height of 139 mm from the base of the cantilever beam. These modifications were implemented to optimize the system for phantom-based experiments while maintaining the high sensitivity and accuracy demonstrated in the original *in vivo* setup.

**Figure 4.**
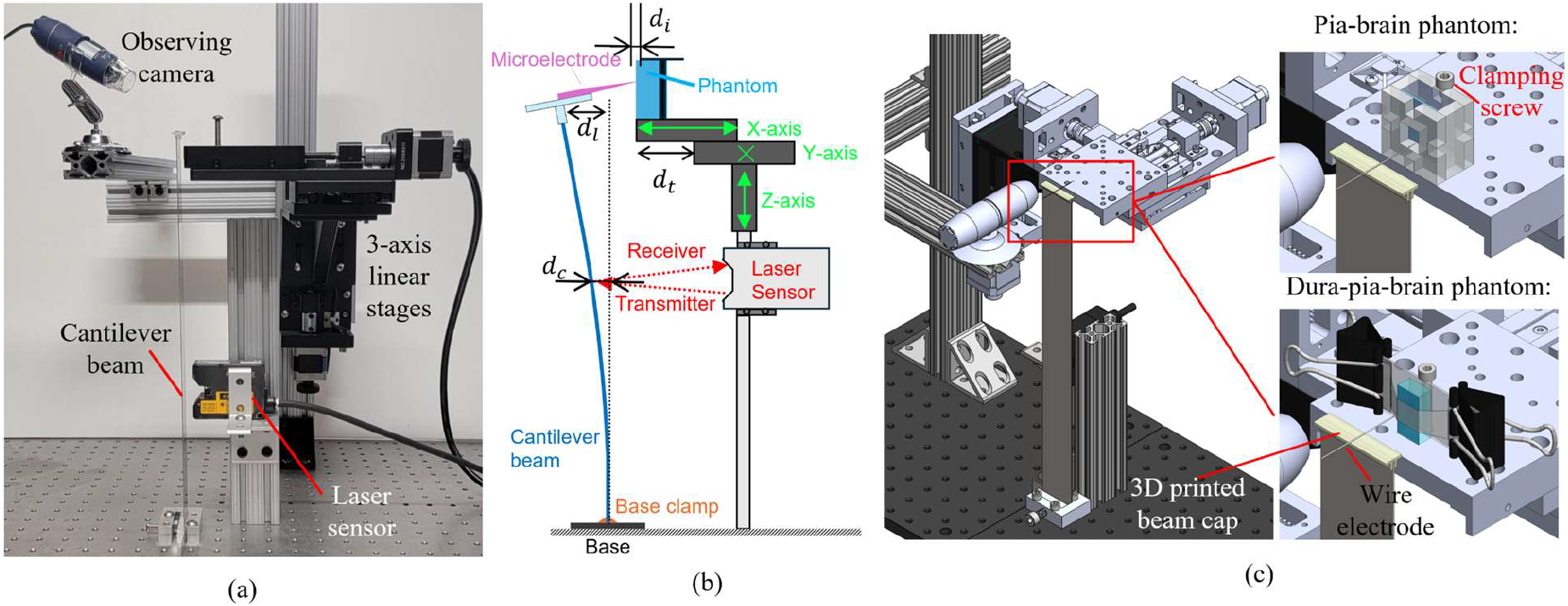
**Overview of the experimental setup for brain-mimicking phantom evaluation: (a) the modified cantilever-beam based force measurement setup for phantom insertion test, (b) illustration of key measurement parameters during insertion for rupture force and dimpling depth calculation, including dimpling depth (***d*_*i*_**), stage translation (***d*_*t*_**), wire tip deflection (***d*_*l*_**) and cantilever beam deflection measured by the laser sensor (***d*_*c*_**), (c) demonstration of the phantom and electrode fixation mechanism during phantom insertion experiments**.

During insertion tests, the phantom assemblies were mounted onto a rigid and precisely aligned support structure. For both the pia-cortex and dura-pia-cortex phantoms, the assemblies were securely positioned on a glass substrate attached to the X-axis linear stage of the force measurement system by a clamping screw (Fig. 4(c)). The 3D-printed top plate, which was fabricated via stereolithography resin printing (Form 3B by Formlabs Inc, Somerville, MA, USA) and mounted on the top of the cantilever beam, held the microwire or silicon probe shank(s) in place during insertion. The microwire or probe shank(s) was secured inside a capillary tube (0.3 mm inner diameter, 0.1 mm wall thickness) with the tip extending beyond the tube for 1.5 mm. The tube was fixed onto the top plate, ensuring stability and balance during testing.

Before each insertion experiment, the laser displacement sensor was calibrated at the cantilever beam’s initial, undeflected state. The phantom was then aligned manually such that the microwire tip was centered and positioned just above the phantom surface without making contact. A live microscope feed provided visual confirmation of this alignment. The insertion was carried out by initiating a constant-speed translation of the X-axis stage at 100 μm/s, a value chosen to replicate *in vivo* insertion speeds used in prior studies for proper validation comparisons. All experiments were conducted at room temperature (25°C), and the phantoms were stored in the refrigerator for less than 24 hours before being used for tests. Also, all phantoms were tested within the same time interval after removal from refrigeration to ensure thermal equilibrium and consistent material properties. Each phantom assembly was discarded after a single insertion to eliminate potential mechanical degradation or residual deformation that could affect subsequent trials. Prior to the insertion of each new microelectrode, the electrode tip was examined under the microscope to confirm integrity and alignment. If the tip was damaged, a new electrode with similar tip geometry would be used instead.

### 2.3. Design of experiment for brain phantom validation

To compare the phantom rupture and dimpling performance as in vivo brain, the validation experiments follow the same insertion device, insertion protocols, and data processing mechanism as the in vivo study for each phantom insertion trial. The primary metrics used for performance validation were the membrane rupture force and the dimpling depth at rupture.

Eleven types of microwire electrodes, same as the in vivo study, were used to test the developed brain phantom. As shown in Fig. 5, they were composed of four different diameters (12, 25, 50, and 100 µm), two different materials (tungsten (W) and stainless steel (SS)), and two different tip geometries (blunt and conical sharp tips). To standardize tip geometry across trials and enable meaningful comparisons, all microwires were pre-characterized using scanning electron microscopy (SEM). Sharpened tips were fabricated through controlled electrochemical etching using a 0.1 M NaOH solution and a constant direct current bias, yielding a smooth conical taper. Blunt tips were prepared through a standard polishing protocol. Briefly, bundles of wires are inserted into capillary tubes. All the bundles are aligned in a 3D printed custom fixture which can fit on the semi-automatic grinder polisher (AutoMet 250 by Buehler Ltd, Lake Bluff, IL, USA). All the bundles inside the capillary tubes are filled with wax (Apiezon Wax X by M&I Materials Ltd, UK). Polishing pads of different grit levels (from 240 to 1200) were utilized along various steps of processing until the wire tips were fully polished. Then the residual wax was carefully removed leaving only the blunt wires ready for use. Representative SEM images of each microwire type are provided in Fig. 5, highlighting tip geometry, surface finish, and diameter consistency. These morphological features were carefully maintained across replicates within each wire group to minimize geometric variability as a confounding factor. The insertion trials were grouped and analyzed based on these wire classifications.

**Figure 5.**
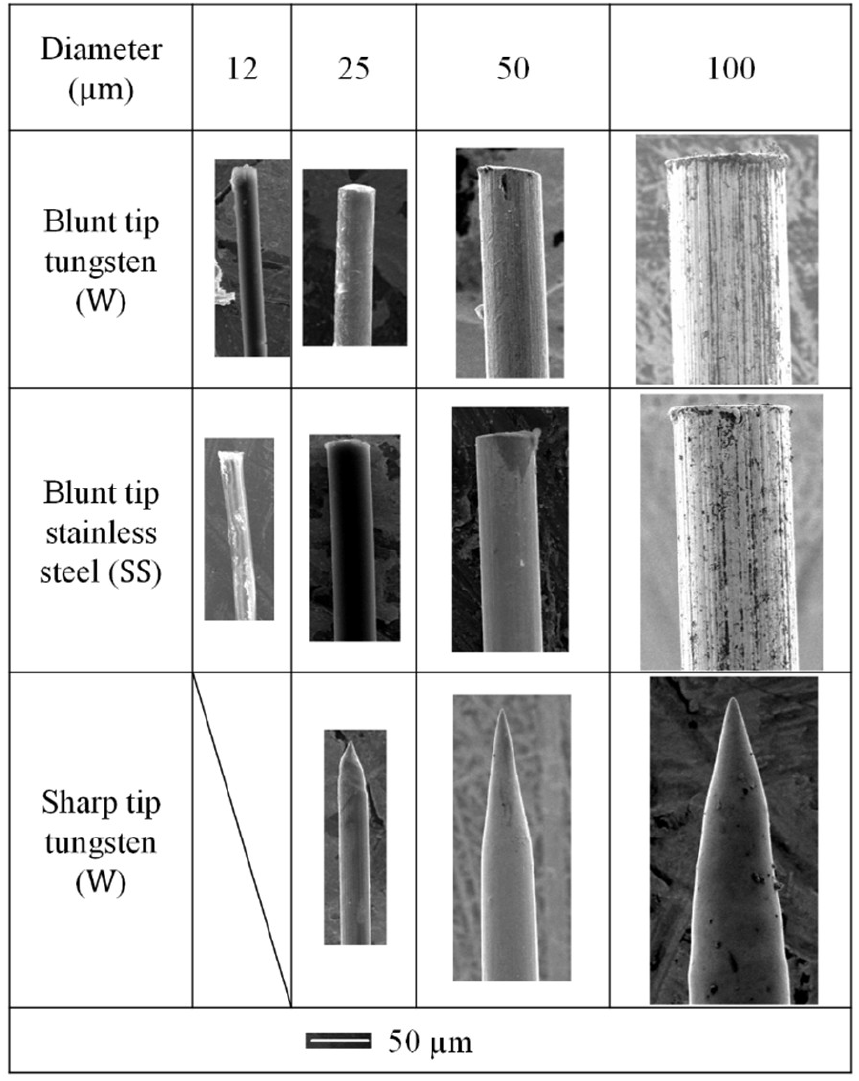
Scanning electron microscopy (SEM) images of representative microwires used in the phantom insertion experiments.

The 12 µm microwires were tested against the pia-cortex phantom only and all remaining microwires were inserted into both the pia-cortex and dura-pia-cortex phantoms for validation measurements. For each condition, at least three repeated trials were performed and a new phantom were used for each test. The microwire tip was examined under microscope to verify the integrity of its geometry. If any deformation or fracture of the tip was observed, the wire was replaced with another comparable tip geometry to maintain uniform tip characteristics across trials. In all experiments, microwire was fed into the phantom until full membrane penetration was confirmed visually via the USB camera feeding and the laser sensor displacement drop. Insertion data were captured continuously throughout the process, from initial contact with the phantom surface until post-rupture relaxation. Trials with visual evidence of microwire misalignment, broken wires, laterally shifted dura layer phantom, or other setup instability were discarded and excluded from the analysis. A new trial would be re-conducted to fulfill the data collection needs under that cutting condition.

Besides microwires, silicon probe shanks were also used for the brain phantom validation against in vivo measurements. A commercial 8-shank silicon array (Buzsaki64 – H64LP probe by Neuronexus, Ann Arbor, MI) were segmented into pieces with one, two, or four parallel shanks connected. Each individual shank was 50 μm wide and 15 μm thick and distances between adjacent shanks in a dual or quadro-shank pieces was 200 μm. These probe shank segments were mounted onto the same glass capillary tube and inserted into pia-only phantom assemblies under identical experimental conditions to those used for microwire testing. Two insertion trials were conducted per shank configuration, yielding a total of six tests.

The data sampling and collection methodology used in this study follows the approach established by the in vivo study [5]. The cantilever beam used in the experimental setup (Fig. 4) was calibrated via both edge pushing experiments and finite element modelling to correlate the beam deflection with insertion force. The linear stage movement along the X-axis (*d*_t_) (Fig. 4) was calculated by multiplying the constant feed rate (100 μm/s) by the total insertion time. Finally, the values of *d*_l_ and *d*_t_ were used to compute the phantom surface displacement (*d*), representing the dimpling depth on the phantom surface at penetration point.

## 3. Results

### 3.1. Multi-layer brain-mimicking phantom parameters

Geometries and material formulas for each brain layer phantom went through iterative tests and tuning via comparison with the in vivo measurement results. The final phantom parameters were as elaborated in the fabrication protocols in Section 2 and key tuning procedures include:

- Thickness of all three phantom layers: The pia and cortex layer thicknesses were tuned by adjusting the pipetting amount of agarose solution based on the silicone cube cross sectional area. The dura layer thickness was mainly tuned via changing the press mold cover slip thickness. The final thickness of the dura, pia, and cortex layers were about 0.15 mm, 0.6 mm, and 9.9 mm accordingly. A thick cortex layer minimized impact of the rigid backend support (acrylic piece or glass slide) on the insertion boundary conditions.
- Material properties of all three phantom layers: The pia and cortex layers were controlled via the agarose solution weight by volume ratio (0.5% for cortex and 1,01% for the pia mater finally). The dura layer property was mainly tuned via the PVC and plastic softener ratio (2:1 for the rat brain match).
- Layer interactions: Two-step Agar curing protocol ensured tight connection between the pia and cortex layers of phantom. The contact between the dura mater phantom and the pia-cortex part was tuned via the length pre-stretch amount (25% in this study’s final finding).

### 3.2. Rupture force and dimpling depth comparison

Figure 6 summarizes the in vivo rupture force and dimpling depth results as reported in previous study [5]. Key findings from the in vivo measurement included that (1) both pia-only and dura-pia penetration showed rupture force and membrane dimpling depth at rupture to be linearly related to the wire diameter, (2) dura-pia rupture force was roughly an order of magnitude higher than the pia-only rupture counterpart, (3) wire sharpening and diameter reduction significantly reduces both rupture force and dimpling depth, and (4) materials does not significantly impact the rupture force or dimpling depth.

**Figure 6.**
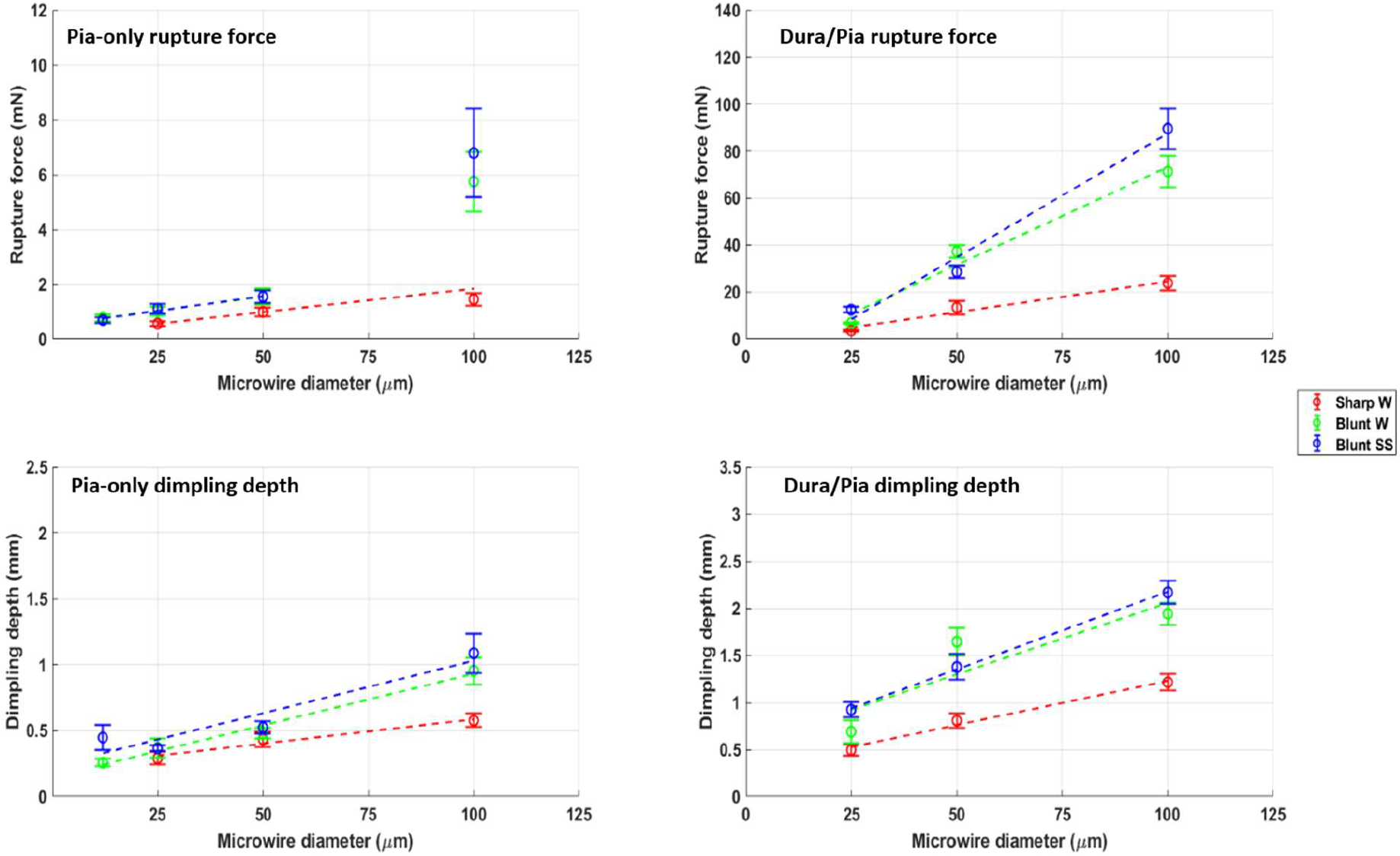
Summary of average rupture force and dimpling depth under all microwire insertion conditions during in vivo insertion tests on Sprague-Dawley rat brain [5]. Error bars present standard errors.

To validate the developed brain-mimicking phantom’s performance, Figure 7 presents the rupture force and dimpling depth results from same wires and same cutting conditions under the same scales. In pia-only insertions, rupture forces remained below 10 mN and dimpling depths under 1.5 mm across all cases, while the dura–pia configuration exhibited significantly elevated rupture forces and dimpling depths. These trends closely paralleled in vivo observations, with linear regression analyses revealing strong correlations between microwire diameter and both rupture force and dimpling depth. The phantom data successfully preserved the relative scaling behavior across tip geometries—i.e., increasing rupture force and dimpling with larger diameters, and decreasing values for sharper tips—thereby validating the phantom’s ability to replicate core insertion mechanics. Notably, the phantom exhibited substantially reduced variability in both rupture force and dimpling depth, as compared to *in vivo* data. These results underscore the phantom’s greatly improved experimental consistency and its utility as a repeatable testing platform for novel neural interface development and insertion mechanics studies.

**Figure 7.**
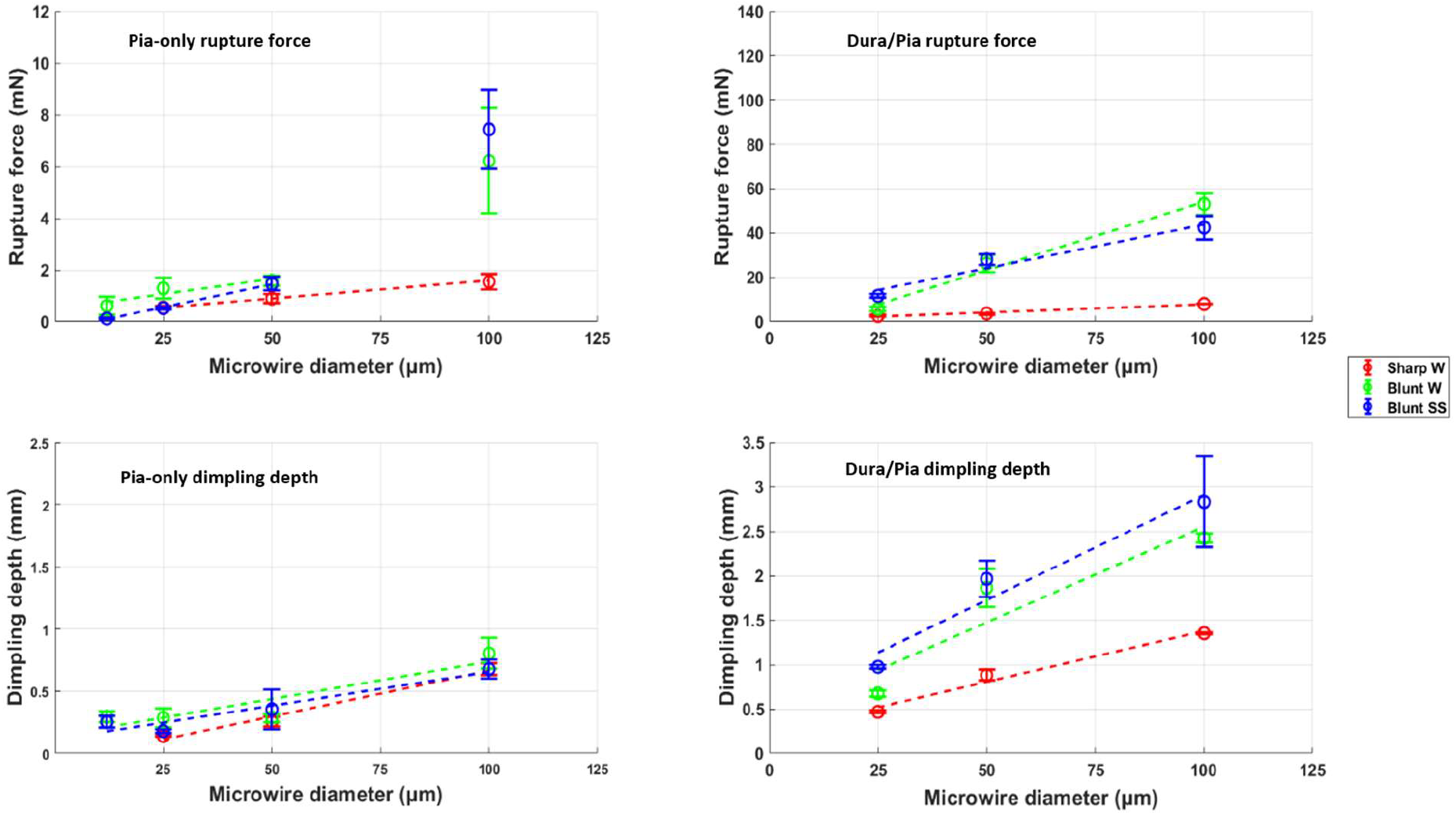
Summary of average rupture force and dimpling depth under all microwire insertion conditions during insertion tests on developed brain phantoms [5]. Error bars present standard errors.

For pia-only insertions, the slope and intercept of the regression lines closely matched *in vivo* values, confirming that the phantom effectively reproduces the force–displacement dynamics at the tissue-electrode interface. Minor deviations were observed at larger diameter wires (100 µm), where the phantom responses showed slightly attenuated increases in rupture force and dimpling depth, likely due to the phantom’s inability to fully replicate the nonlinear viscoelastic properties and anatomical heterogeneity of biological brain tissue and membrane structure. In the dura–pia condition, regression slopes and average rupture force and dimpling depth values were lower than *in vivo*, but overall trends remained consistent. This suggests that while the developed phantom simplified the mechanical complexity of *in vivo* tissue, it remains a valid and robust surrogate for studying insertion behavior and for preliminary testing of novel neural interfaces.

To further investigate the variances on average rupture force and dimpling depth results between in vivo and phantom tests, blunt stainless steel microwire results with comparatively large average value variance were taken as an example and all raw data points from in vivo and phantom trials are plotted in Fig. 8 with horizontal axis showing the rupture force and vertical axis representing the dimpling depth at rupture. As seen, across all conditions, despite variance between average values, phantom data fell within the *in vivo* distribution boundaries, demonstrating close agreement in both magnitude and trend. Notably, phantom results consistently showed reduced variability between repeated trials, reflecting the benefits of the developed brain-mimicking phantom as a low-cost repeatable test platform.

**Figure 8.**
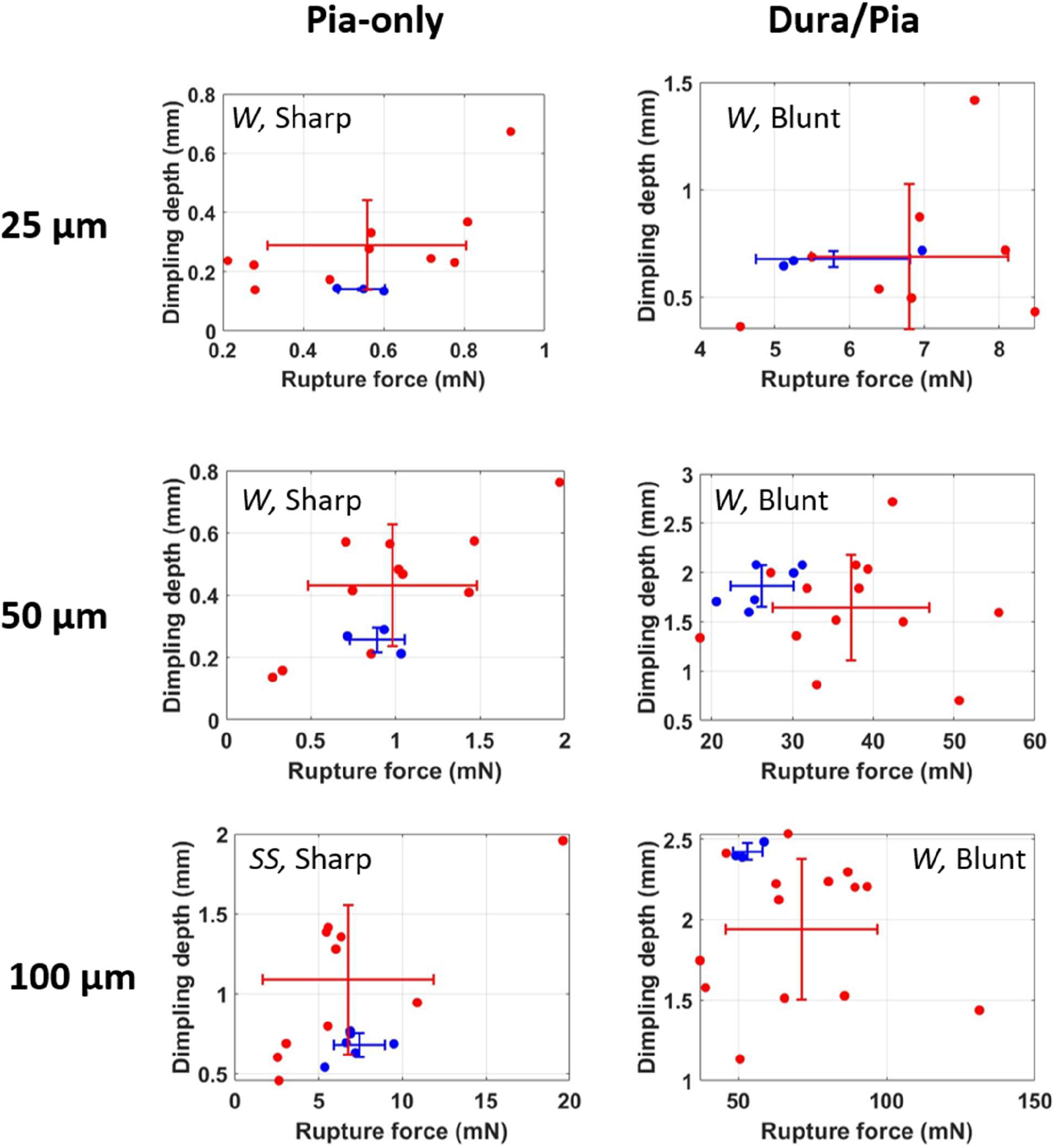
Comparison of rupture force and dimpling depth raw data between phantom (blue) and in vivo (red) tests with stainless steel (SS) and tungsten (W) microwires. Error bars show standard errors for each outcome.

The match between developed brain phantom and in vivo results also showed for silicon probe shank insertion tests, as shown in Fig. 9. For single-, dual-, and four-shank insertion configurations, both rupture force and dimpling depth results from phantom insertion tests showed consistency with in vivo observations, both in linear trends and in magnitudes. Despite the limited sample size (n=2 per group), these findings demonstrate the developed phantom’s capability of replicating planar neural probe insertion performances.

**Figure 9:**
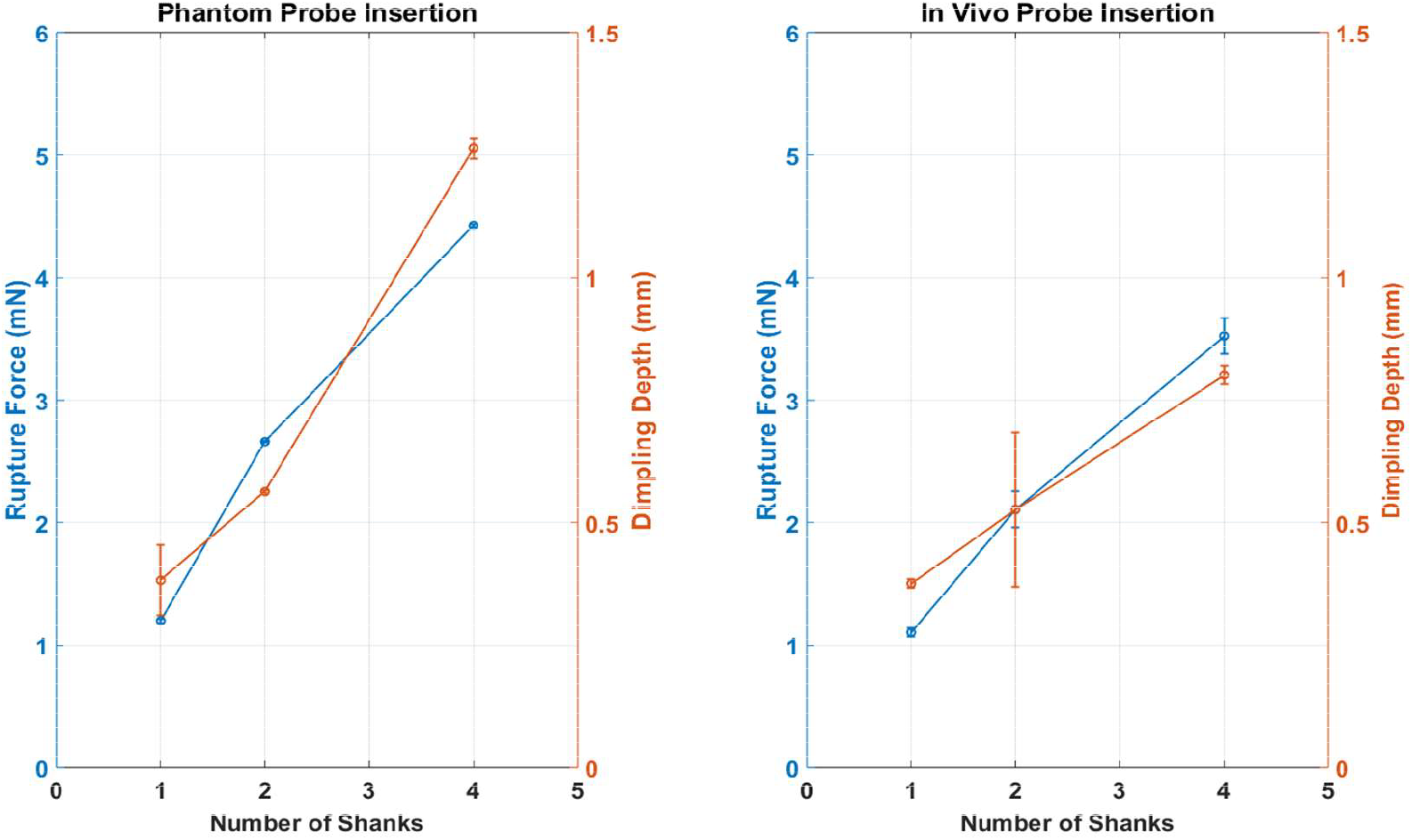
Comparison of rupture force and dimpling depth across single-, dual-, and four-shank silicon probe insertions in phantom (left) and in vivo (right) tests.

**Figure 10.**
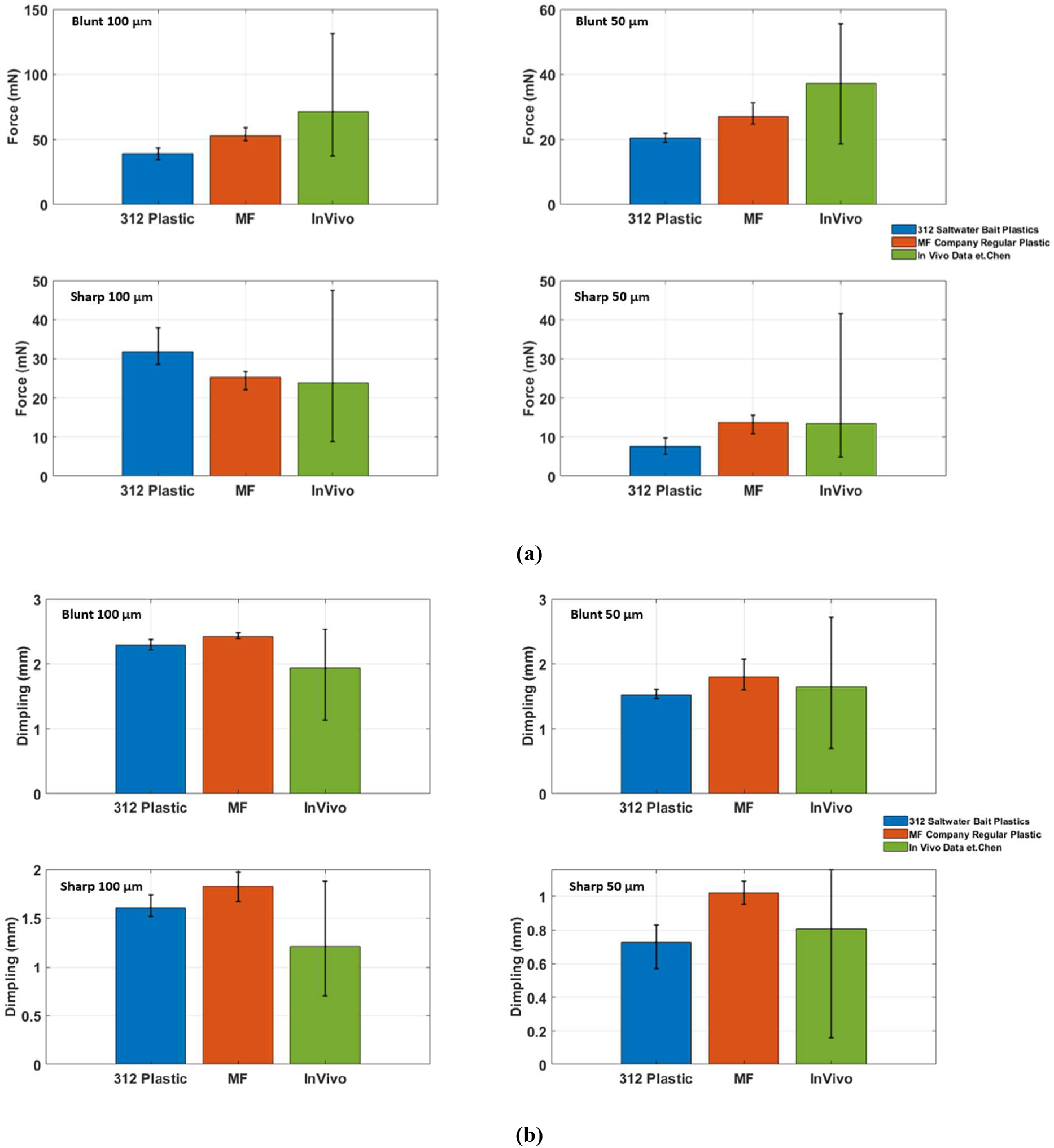
**Comparison of (a) rupture force and (b) dimpling depth across multi-layer phantoms with dura-mimicking layers fabricated using different plastisol materials. Bar plots show average rupture force (mN) or dimpling depth (mm) ± standard deviation for four insertion conditions: 100 µm blunt, 100 µm sharp, 50 µm blunt, and 50 µm sharp microwires. Three groups are presented for each condition: 312 Saltwater PVC plastisol (blue), M&F Manufacturing plastisol (orange), and in vivo rat dura measurements (green)**.

## 4. Discussion

By leveraging in vivo insertion data and the cantilever-beam-based force measurement system, we successfully developed a multi-layer brain-mimicking phantom that replicates in vivo rat brain rupture force and dimpling depth during neural interface implantation process. While in vivo test results showed large variance likely due to tissue condition, animal-to-animal differences, and the complexity of surgical preparation, the phantom instead provided consistent rupture performance within the in vivo dataset range. As a result, the developed phantom could serve as a repeatable insertion test platform for engineering development of new neural interface geometries, materials, and sizes and foster lower-cost and higher-efficiency iterative trials for design optimization. Also, easily accessible insertion tests, including both pia-only and dura-pia insertion scenarios, could enable more comprehensive cutting mechanics studies to generate new understanding of the neural interface insertion process, which would guide further design and development efforts.

While the brain-mimicking phantom formula developed in this study only tried to replicate the Sprague-Dawley rat brain behavior, the overall multi-layer brain mimicking phantom structure could easily be further tuned to fit other animal models and insertion scenarios. The thickness and property of each phantom layer as well as the interlayer bonding could all be independently adjusted, as elaborated in Section 3.1, based on specific targeted animal model needs. Over 600 experimental insertion trials were conducted in this study to finalize the developed brain phantom formula and fabrication protocol. Based on our iterative observations, the following guidelines could be provided for further tuning of the multi-layer brain phantom:

- The soft cortex layer dominates the overall phantom’s dimpling performance. The standard 0.5% agar solution in DI water was used in this study. It was found in our iterative trials that agar concentration below 0.5% would create a phantom that lacked structure integrity, causing it to become fluid and unable to maintain its shape. In contrast, a higher agar concentration could potentially lead to quicker curing and pipette clogging if dispensing large amount. Also, higher concentration agar blocks could yield a brittle material which fractures upon membrane layer penetration and produces inconsistent rupture force and dimpling depth measurement for the whole phantom. Thus, it would be suggested to keep the brain tissue phantom formula close to 0.5% Agar and adjust the dimpling property by changing the overall sample size and modifying the boundary conditions to enable larger or smaller dimpling response.
- In contrast, the pia layer phantom, thanks to the smaller volume and greater baseline concentration, showed greater stability in the presence of material compositional change. Our trials showed that for pia-only insertions, pia formulation of 0.84% to 1.01% agar showed little difference in displaying performance comparable to in vivo data. However, when dura layer was added, only the 1.01% agar kept consistent result as in vivo while the lower concentration ones showed lower rupture force and dimpling depth than in vivo. It would be recommended to take the dura layer contact into consideration before finalizing the pia mater formula.
- The dura mater layer played the biggest role in the mechanical behavior of the phantom because of its specific material properties and stretching ratio. An increase of the plastic component in the mixture yielded a stiffer dura layer, which was associated with greater rupture forces and a slight rise in dimpling depth. Conversely, the increase of the softener ratio—or the decrease in PVC plastic content—resulted in considerably lower rupture forces and a slight reduction in dimpling. Also note that an excess of plasticizer led to brittleness and tearing during insertion, while insufficient plasticizer caused poor material homogeneity and the formation of air bubbles—both of which rendered the layer unsuitable for insertion testing.
- Variations in the original dimensions and elongation ratio of the dura layer phantom PVC film were found also to significantly impact both rupture force and dimpling depth. Specifically, using a larger pre-cut PVC sheet or applying a lower stretch ratio resulted in a thicker dura layer, thereby increasing mechanical resistance to penetration. Determination of the final stretched PVC length also needs to take the pia-brain assembly size into consideration.

PVC plastisol polymer is commonly used for various molding and coating applications. Unlike agarose, it usually has slightly different formula and composition from different vendors. To address potential discontinuation of M&F Manufacturing’s PVC plastisol product, we evaluated alternative formulations for the dura-mimicking layer that maintain comparable mechanical fidelity. A promising substitute was identified in the 312 Saltwater PVC plastisol (sample version), commercially sourced from Bait Plastics LLC (Desloge, MO, USA). This material, mixed at a 2:1 weight ratio with Bait Plastisol Softener—identical to the original formulation—exhibited similar flexibility and puncture resistance to the previously used M&F Manufacturing plastisol. Figure 11 compares the rupture force and dimpling depth across three groups: (i) dura layers fabricated using 312 Saltwater plastisol, (ii) M&F plastisol, and (iii) *in vivo* measurements reported previously. Across different wire diameters (50 µm, 100 µm) and tip geometries (blunt, sharp), the 312 Saltwater plastisol by Bait Plastics LLC (Desloge, MO, USA) produced rupture force and dimpling values that were statistically comparable to those of the M&F Manufacturing plastisol, and within the measured range *in vivo*. In many cases, particularly for sharp microwires—the 312 plastisol samples demonstrated slightly lower variability and comparable or improved matching to *in vivo* dimpling depths.

While the developed brain mimicking phantom showed promising comparison results against the in vivo data, it would still deviate from the real tissue performance of certain cutting conditions change. The phantom does not fully replicate the nonlinear viscoelastic response of real brain membranes and tissues, and does not include considerations of complicated brain features like fibrous structures of the membrane and arteries within the brain. Also, the phantom developed in this study mainly focused on the rupture and dimpling properties of the brain membranes and tissue. Further development will be needed if other electrical, optical, and chemical properties are of interest. Thus, the developed phantom will be beneficial for benchtop insertion tests but ultimate evaluation of the overall neural interface performance will still be based on in vivo test results.

## Conclusion

This study elaborates the design, fabrication, and validation of a multi-layer, brain-mimicking phantom tailored for replicating the rupture force and dimpling depth of the brain during neural interface insertions through the brain membrane layers (both pia-only and dura-pia penetrations). The developed multi-layer phantom is composed of agarose-based pia mater and brain tissue mimicking layers and a PVC-based dura mater mimicking layer pre-stretched and wrapped around the pia-brain assembly. All three layers are fabricated using easily accessible materials and tools following scalable and replicable protocols. Phantom insertion tests using various types of microwires and silicon probe shanks delivered comparable results as previous in vivo Sprague-Dawley rat brain test measurements, but with largely reduced variability between repeated trials, indicating validation of the developed phantom and great potential of using it as a reliable benchtop insertion test platform for novel neural interface development, surgical technique refinement, and fundamental insertion mechanism studies. Beyond the specific Sparague-Dawley formula developed, the modular design of the multi-layer phantom also enables further tuning of different layers’ properties towards other animal models, neural interface designs, and implantation conditions.

## 6. Acknowledgement

The authors acknowledge the financial support from the National Institutes of Health (NIH) under award number R15NS133861 and the National Science Foundation under award number 2347299.

## 7. Conflict of Interest

There are no conflicts of interest.

## Notes

### Competing Interest Statement

The authors have declared no competing interest.

